# Phosphorylation of KRP3, a KIP-related protein by MPK3 modulates rice tiller and seed number by fine-tuning cell division

**DOI:** 10.1101/2024.04.18.590116

**Authors:** Gopal Banerjee, Sarvesh Jonwal, Uttam Pal, Balakrishnan Rengasamy, Dhanraj Singh, Alok Krishna Sinha

## Abstract

During the cell cycle process, multiple inhibitors function as checkpoint regulators ensuring an impeccable duplication of genetic material. Kip-Related Proteins (KRPs) are a group of plant-specific cell cycle inhibitors and are known to regulate plant architecture and yield by differentially regulating cell division. KRPs, though functionally conserved, are dynamic in their amino acid composition. Interestingly, some KRPs are specific to dicots, and some are for monocots. In this study, we have identified that the presence of KRP3 and KRP6 is specific to the Poaceae family of monocots. An in-depth study of *KRP3* showed a strict regulation of its expression in actively dividing cells during the G1-S phase progression of cell division. Our study identified KRP3 as a phosphorylation target of MPK3. The phosphorylation enhances KRP3 protein stability, which leads to strong inhibition of cell proliferation. By generating knock-out lines of *krp3*, *mpk3* and double knock-out *krp3mpk3*, our study demonstrated that the MPK3-KRP3 module regulates rice root and shoot development as well as tiller and seed numbers. It was observed that KRP3 regulate the rate of cell division in a dose-dependent manner. The study, in a nutshell, establishes the role of KRP3 as an important regulator of rice vigor and yield.

## Introduction

The synchronized orchestra of cell cycle regulation is governed by multiple groups of proteins. In land plants, two groups involved in cell cycle checkpoint regulation are known as Inhibitor of Cyclin Dependent Kinase (ICK)/Kip Related Protein (KRP) and Siamese/Siamese-Related Protein (SMR). KRPs do not show much sequence similarity with their mammalian counterparts except for a conserved CDK and Cyc binding motif at their C-terminal region. SMRs, on the other hand, are completely plant-specific cell cycle inhibitors. These two distinct groups of inhibitors also function distinctly in regulating the cell cycle, where KRPs can inhibit cell division at both G1-S and G2-M checkpoints, while SMRs are limited only in the G2-M phase regulation. Irrespective of their functional similarities, KRPs can be grouped into three distinct groups depending on their sequence similarity. The KRPs, with their vast number, are responsible for maintaining distinct plant architecture to stress responses. Expression of *KRP1* under CaMV35S promoter reduces endoreduplication and induces cell death in *Arabidopsis* leaf trichomes (Schnittger et al., 2003). Overexpression of *KRP2* in *Arabidopsis* leads to reduced lateral root number. *KRP1* and *KRP2* overexpression also affects root-knot nematode offspring number and gall size. Studies also found that the protein level of AtKRP4 determines S phase initiation time in shoot stem cells (D’Ario et al., 2021). Unlike other KRPs, KRP6 showed an antagonistic effect in cell cycle regulation as *KRP6* overexpression promotes cell division rate in *Arabidopsis*. Mutants of *krp5* showed shorter root lengths, as KRP5 shows strong expression in root apical meristem and is responsible for root cell elongation (Wen et al., 2013). It was only when a quintuple KRP mutant was generated and studied, that an increase in organ and seed size was observed resulting from an increased cell proliferation rate (Cheng et al., 2013). In the monocot model system, rice, the study of *KRP1* overexpression showed longer cell size in the leaf along with small seed size (Barrôco et al., 2006). Interestingly, another study of *krp2* mutant along with *krp1krp2* double mutation also showed reduced grain filling, leading to smaller grain size as well as a low germination rate (Ajadi et al., 2020). A lower seed setting rate was also observed in rice *krp4* and *krp5* mutants (Xu et al., 2023).

For keeping the cell division on the go, the protein levels of KRPs at different cell cycle stages are strictly regulated, predominantly by ubiquitination-mediated proteasomal degradation. FBL17, an F-Box-like protein, has been found to interact with multiple KRPs like KRP4, KRP6, and KRP7 and mediate their degradation (D’Ario et al., 2021; Pan et al., 2021). Other than FBL17, KRP1 alone have been identified to be a target of other E3 ligases likely SCF^SKP2b^, RING protein RPK along with UBL/UBA protein RAD23B (Li et al., 2020). In the monocot system, maintenance of KRP protein level is mostly unknown except for a recent report of F-box protein 3 (*FB3*) mediated proteolysis of KRP4 and KRP5 in rice (Xu et al., 2023).

Like cell division, Mitogen-Activated Protein Kinase (MAPK) signaling is a conserved signaling cascade involved in regulating diverse pathways associated with organ development to stress responses in both plants as well as animal systems (Jonwal et al., 2022; Manna et al., 2023; Mittal et al., 2022). *AtMPK3* and *AtMPK6*, the two most studied MAP kinases from *Arabidopsis*, are known to regulate ovule and stomata development, hypocotyl length and microRNA biogenesis (Bhagat et al., 2022b; Jiang et al., 2022; Sethi et al., 2014). *YODA* and *MPK6* regulate root development by influencing auxin level and cell division plane orientation (Smékalová et al., 2014). Another signaling cascade consisting of MKK6 and MPK13 regulates lateral root development in *Arabidopsis* (Zeng et al., 2011). In rice, a OsMKKK10-OsMKK4-OsMPK6 module regulates grain size and number as well as leaf angle by targeting WRKY53 as a BR response signal (Tian et al., 2021). This pathway is negatively regulated by Grain Size and Number 1 (GSN1) / MAP Kinase Phosphatase 1 (MKP1). It was also observed that the larger seed size in *gsn1* is due to a high cell division rate with no effect on cell elongation (Guo et al., 2018). Though MAP Kinase-mediated cell cycle alteration has been observed but no direct target of cell cycle protein was known in plants until recently, where WEE1 and SMR1 are phosphorylated by MPK3 and MPK6 under UV-B exposure (Banerjee et al., 2023). Another report suggests that MPK3, MPK4 and MPK6 target the E2F family transcription factor, E2F2 and negatively regulate its function (Singh et al., 2023).

Yield potential and quality are one of the most important criteria for cereal crops. In grain crops, yield is directly associated with seed per plant and seed weight. The latter is again determined by the seed size (Li et al., 2021). Yield is a multi-locus regulated trait, and cell division regulators are a major player in them (Burgess et al., 2023). Cell division can also regulate crop yield indirectly by many other aspects like by regulating organs like leaf and root size and structure (Burgess et al., 2023).

In this study, we have identified KRP3, a unique KRP present only in the Poaceae family of monocot lineage and is a phosphorylation target of MPK3. CRISPR-Cas9 mediated mutation in *krp3* exhibited an identical trend of plant growth with *KRP3* overexpression plants, like shorter root and internode length as well as a reduced tiller and seed number. It was further observed that irrespective of a cell cycle inhibitor, the presence of KRP3 is very crucial for maintaining cell division rate in actively dividing zones. *mpk3* mutation also depicts similar phenotypes and functions upstream of KRP3 to maintain rice organ size. Further, it was observed that overexpression of *KRP3^EE^* (phospho-mimetic form of KRP3) showed stronger cell division inhibition over *KRP3* or *KRP3^AA^*(KRP3 phospho-dead form) overexpression plant. The investigation further revealed that phosphorylation of KRP3 at S^17^ ^&^ ^82^ positively influence its stability.

## Results

### KRP3 is involved in regulating rice organ size

KRPs have previously been divided into three broad groups depending on their sequence similarity. One group contain KRPs from both monocot and dicot lineage, while two others consist of KRPs solely from the dicot or monocot lineage (Yang et al., 2011). In this study, a detailed phylogenetic tree has been constructed with all KRPs reported from monocot plants (**Sup. Fig. 1**). It was interesting to observe that *KRP1*, *KRP4* and *KRP5* are present consistently over twenty-two genera while *KRP3* and *KRP6*, have been reported in 9 and 12 genera respectively, that belongs specifically to the Poaceae family. To elucidate the functionality of this considerably modern KRP, a knockout and overexpression line of KRP3 was developed. *krp3* knockout mutants were developed using the CRISPR-Cas9 tool (**Sup. Fig. 2A**). Out of all the mutant lines identified for *KRP3*, two independent lines with either single nucleotide insertion (*krp3-1*) or deletion (*krp3-2*) in homozygous condition were selected for the study (**Sup. Fig. 2B, C, D**). Along with *krp3* mutants, transgenic plants overexpressing *KRP3* under CaMV-35S promoter were also generated (**Sup. Fig. 3A**). Three independent *KRP3* overexpression lines showing highest level of *KRP3* expression were selected for this study (**Sup. Fig. 3B**). Plants generated from callus transformed with pCAMBIA1300 vector were used as Vector Control (VC) (**Sup. Fig. 3C**).

A comparative analysis of multiple morphological as well as yield-related traits for the *krp3* mutant and KRP3-OE lines were conducted. Plant height was measured for Tp309, *krp3-1*, *krp3-2*, VC, *KRP3-OE1*, *KRP3-OE2* and *KRP3-OE3* lines. No significant difference was observed between the Tp309 and VC plants. Interestingly both *krp3-1* and *krp3-2* showed 17% and 12% reduction respectively in plant height compared to Tp309 (**Fig. 1A, B**). Similar to *krp3* mutants, *KRP3* overexpressed plants also showed a significant (∼15%) reduction in plant height compared to Tp309 and VC plants (**Fig. 1A, B**). A similar pattern of reduction was observed in the case of tiller number and first internode length. Though *KRP3-OE* lines showed a minor reduction in tiller number (∼10%), a significant reduction was observed in *krp3-1* and *krp3-2* lines (37% and 43% respectively) (**Fig. 1C**). Significant reduction in internode length was observed when the length of first internode of *krp3* mutants (26%) and *KRP3-OE* lines (∼20%) was measured and compared to Tp309 and VC plants (**Fig. 1D**). To determine the tissue-specific expression of *KRP3*, ProKRP3::GUS line along with VC-GUS line were developed (**Sup. Fig. 4A, B**). A strong GUS activity was detected in elongating internodes of ProKRP3::GUS line which was not visible in control plant (VC-GUS) (**Sup. Fig. 5**). When leaf length and width were measured, no apparent effect was observed in *krp3-1* and *krp3-2* leaf compared to Tp309. Though the mutant leaf length was unaffected, the *KRP3-OE* leaf length was significantly smaller (**Fig. 1E, F & G**).

**Figure 1:**
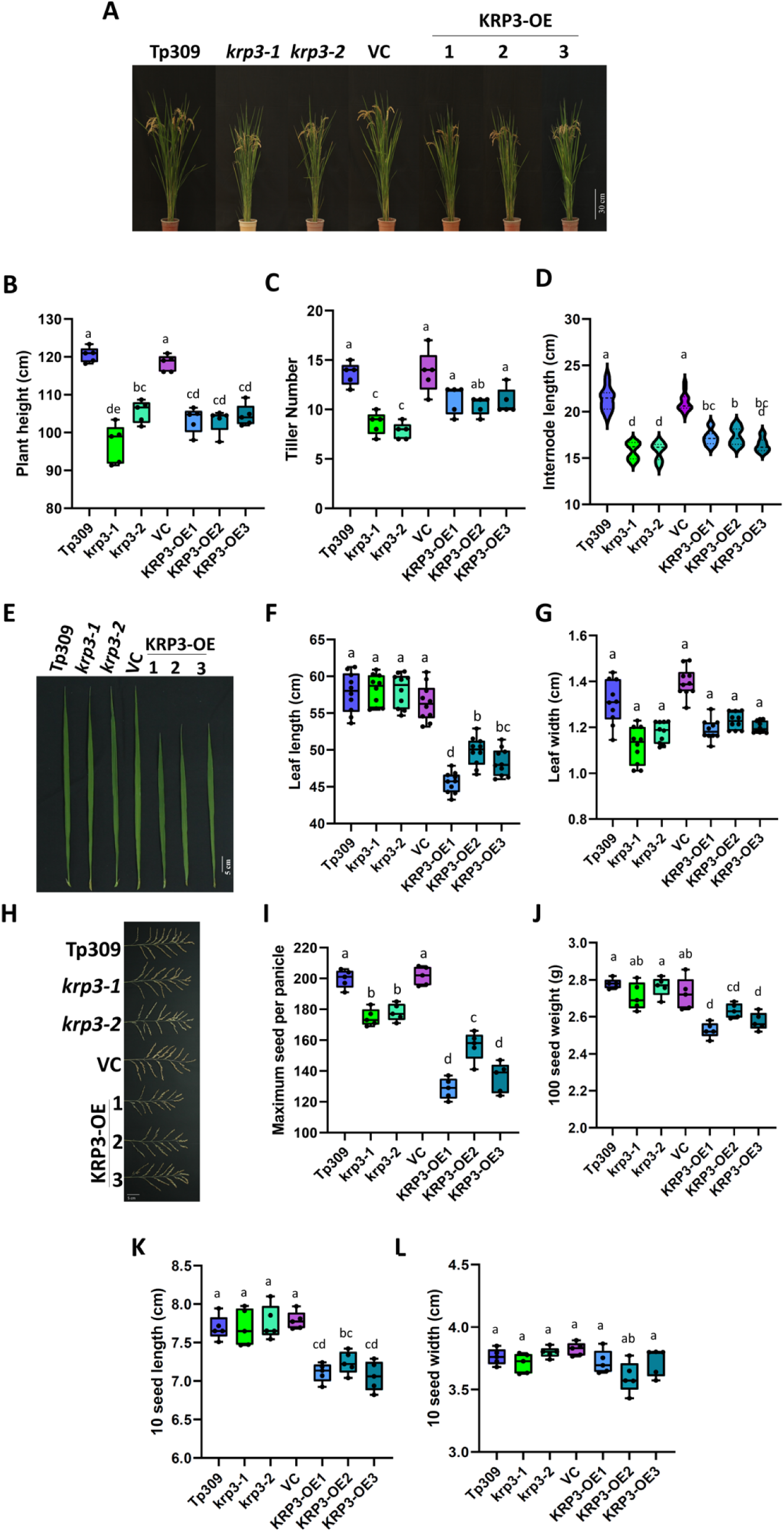
*KRP3* regulates rice vegetative growth and yield. **A.** Pictorial image of Tp309, *krp3-1*, *krp3-2*, VC, KRP3-OE1, KRP3-OE2 and KRP3-OE3 mature plant respectively. **B, C.** Box plot representing the plant height (B) and tiller number (C) of all the lines (n=5). Both *KRP3* mutant and over-expressed plants showed reduced plant height and tiller number. **D.** Violin plot showing 1^st^ internode length of all the lines (n>20). Reduced internode length was observed in mutant and overexpression lines of *KRP3* compared to Tp309 and VC. **E.** Photograph depicting the last mature vegetative leaf of all the lines. **F, G.** Leaf length (F) and width (G) of all the lines (n=10). Reduction in leaf length was found only in KRP3-OE lines while leaf width was unchanged. **G.** Mature panicle of all the lines showing the difference in seed number. **H, I.** Maximum seed number of primary panicle and 100 seed weight of all the lines. Reduction in seed number was evident in *KRP3* mutant and OE lines while reduced seed weight was only observed in KRP3-OE lines. **J, K.** Box plot representing seed length (J) and width (K) of mature seeds (n=10). KRP3-OE lines exhibited only reduced seed length while *krp3-1* and *krp3-2* showed no change in seed size. One-way ANOVA and Tukey’s multiple comparisons test was conducted to compare the differences. Different alphabets represent significant changes observed in Tukey’s multiple comparisons test.

After the vegetative growth parameters, seed traits were evaluated in all the lines. *krp3-1* and *krp3-2* lines exhibited significant reduction (10% and 12% respectively) in maximum seed number in primary panicle compared to Tp309, while the reduction was more pronounced in KRP3-OE lines (35%, 21% and 32% respectively) (**Fig. 1H, I**). Further, precise measurement of seed weight and size revealed no change in *krp3* mutant seeds compared to Tp309 (**Fig. 1J, K & L**). Interestingly, *KRP3-OE1*, *KRP3-OE2* and *KRP3-OE3* also exhibited a reduction in seed weight (6-8%) (**Fig. 1J**). Measurement of seed length and width indicated that seed weight reduction in KRP3-OE lines was primarily due to altered seed length, as *KRP3-OE1*, *KRP3-OE2* and *KRP3-OE3* showed a significant reduction in seed length (6-8%) compared to Tp309 or VC seed while seed width was unaffected (**Fig. 1K, L**). Overall, the data indicated that interfering with the expression of *KRP3* somehow negatively regulates the rice plant’s growth and development.

### KRP3 regulates cell proliferation in rice root

Considering the negative effect of *KRP3* knockout as well as over-expression in mature plants growth, seedling growth rates were analyzed in all the lines (**Fig. 2A**). Shoot and root length of fourteen-day-old *krp3-1* and *krp3-1* seedlings showed retardation in growth like the mature plants, when compared to Tp309 (**Fig. 2B, C**). The overexpression lines of *KRP3* also showed growth retardation comparable to the *krp3* mutants (**Fig. 2B, C**). A strong GUS activity was also detected in the ProKRP3::GUS line, in root tip as well as in emerging leaf (**Sup.** Fig 6). To understand the cause of root growth retardation, cell length as well as width was measured from the maturation zone of root. It was observed that compared to Tp309, both *krp3* mutant and overexpression lines have 1.7 and 1.7-1.3 times longer cell lengths, respectively (**Fig. 2D**). However, cell width was comparable between Tp309 and *krp3-1*, *krp3-2*. It was also observed that cells of KRP3-OE lines were significantly wider, unlike others (**Fig. 2E**). To determine the rate of cell division, EdU incorporation assay was conducted. *krp3* mutant root showed a significantly lower (33-35%) cell proliferation rate compared to Tp309 (**Fig. 2F, G**). A similar reduction in cell division rate was observed in KRP3-OE lines (**Fig. 2F, G**). KRP3 transcript abundance was also studied in Tp309 root using S-phase (Hydroxyurea (HU), Mimosin) and M-phase cell cycle inhibitors (Propyzamide, Oryzalin). Twenty to twenty-five-fold higher KRP3 transcript abundance was observed in S-phase inhibitor-treated roots, while no significant change was observed in M-phase inhibition (**Fig. 2H**). HU sensitivity assay was conducted in the seedlings of all lines to evaluate the role of KRP3 as an S phase checkpoint regulator. *krp3-1* and *krp3-2* mutants exhibited a strong reduction in root growth compared to Tp309 indicating a hypersensitivity towards HU (**Fig. 2I**). Though *krp3* was hypersensitive, root length reduction rate of KRP3-OE lines was similar to that of Tp309 and VC lines (**Fig. 2I**). Cell size analysis of HU treated roots showed that the cell length of Tp309 was higher than that of *krp3* mutants (**Fig. 2J**). Interestingly, the cell length and cell width of HU-treated KRP3-OE lines were similar to that of Tp309 and VC root (**Fig. 2J, K)**. The analysis of different transgenic lines revealed that KRP3 protein content is crucial for cell proliferation. Findings from the HU sensitivity assay also indicate a potential post-translational regulation of the KRP3 protein.

**Figure 2:**
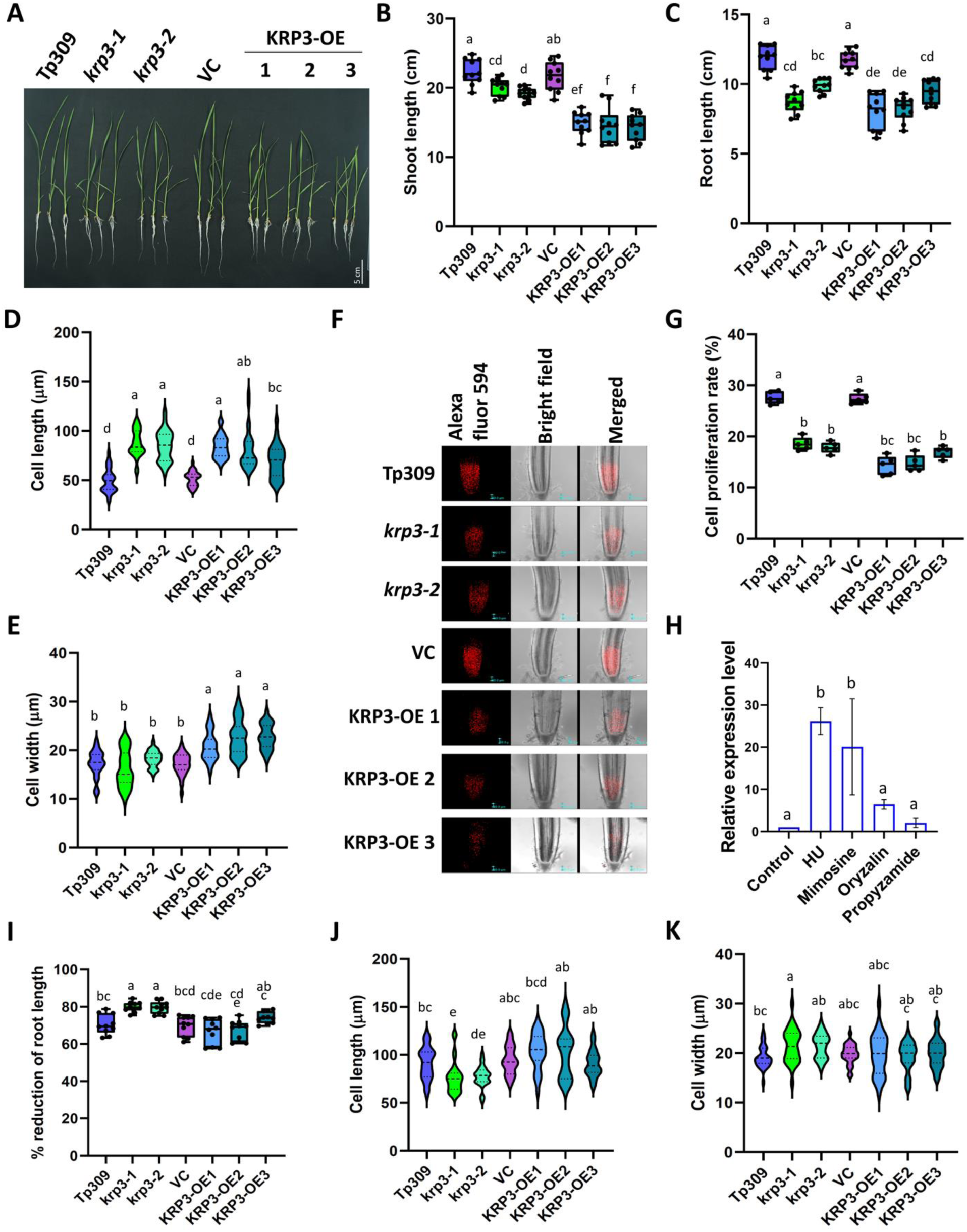
KRP3 maintain cell proliferation by regulating G-S checkpoint. **A.** Image showing fourteen-day-old seedlings of Tp309, *krp3-1*, *krp3-2*, VC, KRP3-OE1, KRP3-OE2 and KRP3-OE3. **B, C.** Shoot (B) and root (C) length of the fourteen-day-old seedlings (n=20). Both *KRP3* mutant and OE lines showed a reduction in shoot and root length compared to controls. **D, E** Violin plot depicting mature root cell length (D) and width (E) of all the lines (n=50). Though cell width was unaffected in *krp3-1* and *krp3-2*, *krp3* knockout as well as overexpression lines showed longer cell length indicating extended cell elongation. **F**. Image of EdU-Alexa fluor 594 labeled rice root observed under confocal microscope. **G.** Rate of cell proliferation (number of cells undergoing S phase) at root division zone. *KRP3* mutant as well as overexpression lines indicated lower cell proliferation compared to Tp309 and VC. **H.** Expression of KRP3 in the presence of S phase (HU, Momosine) and M phase (Propyzamide, Oryzalin) cell cycle inhibitors. Higher KRP3 transcript abundance was observed in the presence of S-phase inhibitors after 4h. **I.** Percent root length reduction in the presence of HU in growth media (n=20). *krp3-1* and *krp3-2* showed hypersensitivity towards HU. **J, K.** Cell length (J) and width (K) of root grown with 2.5 mM HU for seven days (n=50). *krp3-1* and *krp3-2* showed shorter cell length in the presence of HU, while no difference in cell width was observed. One-way ANOVA and Tukey’s multiple comparisons test was conducted to compare the differences. Different alphabets represent significant changes observed in Tukey’s multiple comparisons test. In the case of gene expression analysis, significance was calculated using Dunnett’s multiple comparisons test.

### KRP3 is a phosphorylation target of MPK3

A recent study has shown the involvement of MAP kinases like MPK3, MPK4 and MPK6 in regulating cell division in rice in response to HU-mediated cell cycle inhibition (Singh et al., 2023). Also, our analysis of *krp3* mutant and KRP3-OE lines indicated that there may be a post-translational modification of the KRP3 protein in regulating the cell cycle. To elucidate whether MAP kinase is involved in KRP3 regulation via phosphorylation, the interaction of KRP3 with MAP Kinases was checked. Three rice MAP kinases, namely, MPK3, MPK4 and MPK6 were selected for yeast two-hybrid interaction study with KRP3. The interaction study indicated a specific interaction of only MPK3 with KRP3 and not the other two MAP kinases (**Fig. 3A**). The specificity of the KRP3-MPK3 interaction was further validated using GST pull-down assay. For this assay, GST-tagged KRP3 was used to co-purify His tagged MPK3, MPK4 or MPK6. Western blot of output samples developed using anti-His antibody showed that only MPK3 got co-purified with KRP3 (**Fig. 3B**), validating the findings of the yeast two-hybrid assay. The identified interaction was further verified *in planta* using split luciferase complementation (SLC) and bimolecular fluorescence complementation (BiFC) assays. For the SLC assay, KRP3 was cloned with the C-terminal domain of firefly luciferase, while MPK3 was cloned with the N-terminal domain. Both the constructs were either solely with respective empty vectors or together were co-infiltrated into four different portions of *N. benthamiana* leaf. Luminescence generated by luciferase using luciferin as a substrate was captured using a CCD camera. Luminescence was only observed where both KRP3 and MPK3 were infiltrated together. No luciferase activity was observed in leaf portions where KRP3 or MPK3 were infiltrated with either counter empty vector or both empty vectors (**Fig. 3C**). For BiFC assay, KRP3 cloned with nYFP fraction and MPK3 with cYFP fraction were co-infiltrated in *N. benthamiana* leaf. Both the genes were also co-infiltrated with complimenting empty vectors as control. YFP signal was observed under confocal microscope only when KRP3 and MPK3 were co-infiltrated together (**Fig. 3D**). To determine the KRP3 phosphorylation by MPK3 and the exact site of phosphorylation, *in-vitro* kinase assay was employed. KRP3 was used as the substrate, while MPK3, or MPK3 activated using AtMKK4^DD^, was used as an upstream kinase. Trans-phosphorylation activity of MPK3 activated using AtMKK4^DD^ or without AtMKK4^DD^ activation was confirmed using MBP as a substrate and GST as negative control (**Sup. Fig. 7A**). After confirmation of KRP3 as a phosphorylation target of MPK3, the exact site of phosphorylation was investigated. Four possible phosphorylation sites, S17, S82, S112 and S168 identified *in-silico,* were taken for consideration (**Sup. Fig. 7B**). Four individual constructs were developed where Ser was mutated to Ala in three identified positions. An additional construct where all four Ser residues were converted to Ala was developed (**Fig. 3E**). Bacterially expressed proteins were used for *in-vitro* kinase assay. Phosphor image confirmed that Ser at positions 17 and 82 are the target site for MPK3-mediated KRP3 phosphorylation (**Fig. 3F, Sup. Fig. 7C**). These experiments validated that KRP3 is not only a molecular interacting partner of MPK3 but also a phosphorylation target.

**Figure 3:**
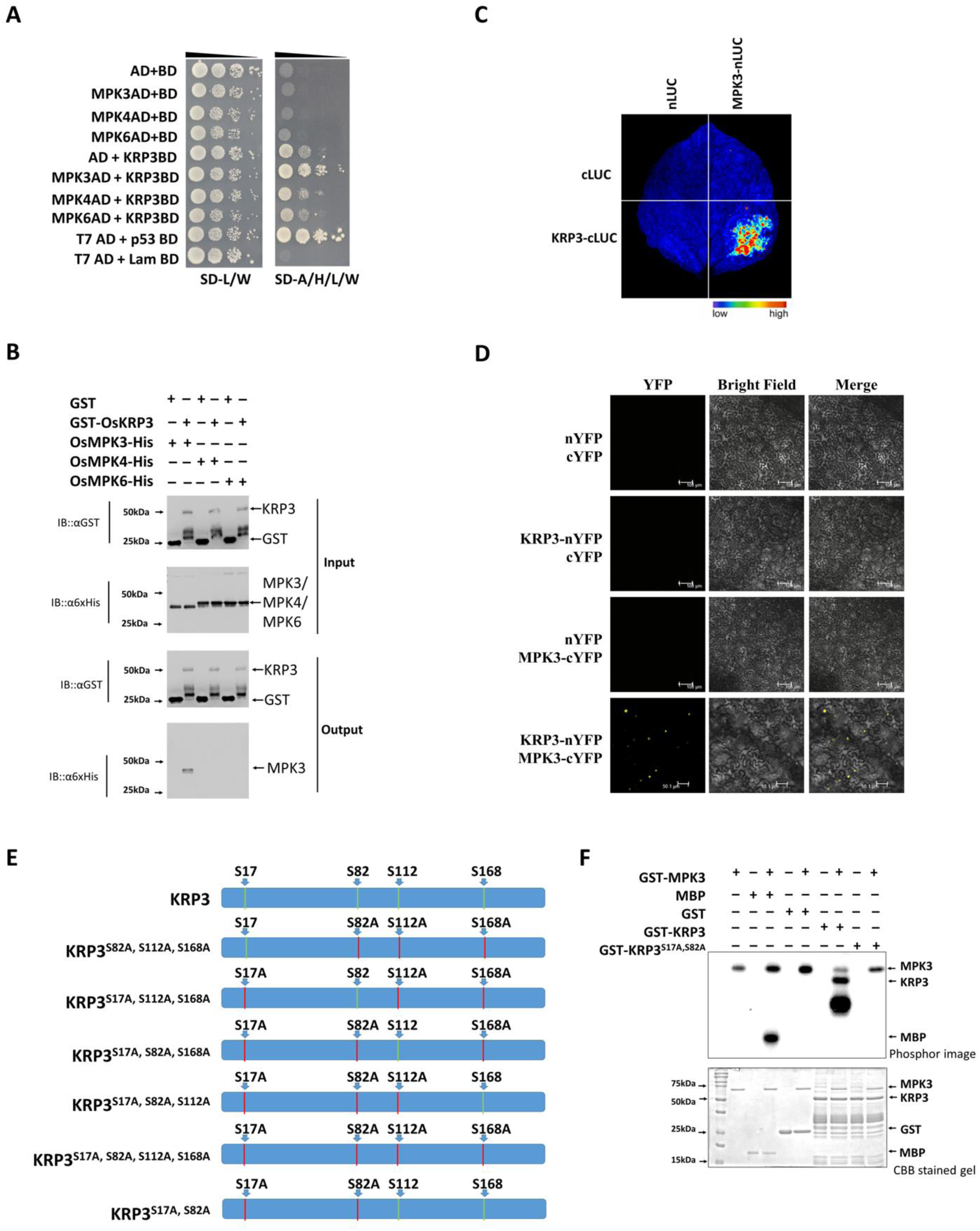
MPK3 interacts and phosphorylates KRP3 at S17 and S82 position. **A.** Yeast two Hybrid interaction assay of KRP3 with MPK3, MPK4 and MPK6. KRP3 was cloned with GAL4 DNA binding domain (BD), while MAP Kinases were cloned with the activation domain. Y2H gold yeast strain was used for this study. Transformed colonies were grown in SD-LW and SD-AHLW supplemented media. Serial dilution of 1, 0.1, 0.01 and 0.001 were grown in supplemented media. A slow growth in KRP3BD-AD transformed colony was observed in SD-AHLW media, while strong growth was observed in MPK3AD-KRP3BD transformed colony. **B.** In-vitro pull-down assay using GST-KRP3 as prey and MAP Kinase as a bait, shows that MPK3 gets co-purified by GST-KRP3, unlike the other two MAP kinases. **C. D.** Split Luciferase Complementation (SLC) assay and Bimolecular Fluorescence Complementation (BiFC) assay demonstrating *in-planta* interaction of KRP3 and MPK3. Positive interaction between KRP3 and MPK3 was detected in SLC as the activity of luciferase was only visible when KRP3 and MPK3 were co-infiltrated (C). Similar observations were made in the BiFC assay, YFP signals were only detected when KRP3 and MPK3 were co-infiltrated (D). **E.** Graphical representation of KRP3 protein structure and position of SP motifs within the protein along with different site-directed mutagenesis construct developed to confirm phosphorylation site of KRP3. **F.** *In-vitro* kinase assay of MPK3 using KRP3 as substrate showing KRP3 gets phosphorylated by MPK3 at Ser 17 and 82 position. MBP was used as positive control and GST as negative control for kinase assay. The kinase assay was repeated three times with similar results.

### KRP3 and MPK3 function in a linear pathway

Once it was established that KRP3 is a phosphorylation target of MPK3, the dynamics of this interaction were checked *in-planta* using *mpk3* as well as *krp3mpk3* double mutant plants. Mutants for MPK3 were developed using CRISPR-Cas9, and two independent lines with single nucleotide addition or deletion were selected for further study (**Sup. Fig. 8 A-D**). For *krp3mpk3* double mutant, a single line with single nucleotide addition in the coding frame of KRP3 and MPK3 was selected (**Sup. Fig. 9 A-B**). *mpk3-1* and *mpk3-2* lines showed moderate reduction (∼6%) in plant height (**Sup. Fig. 10A, B**). Reduction in plant height was also evident in *krp3mpk3* double mutant to that of Tp309 (15%) (**Fig. 4A, B**). A similar trend of reduction in internode length (11 & 8%) as well as tiller number (25%) was observed in *mpk3-1* and *mpk3-2* compared to Tp309 (**Sup. Fig. 10C, D**). Evaluating *krp3mpk3* for internode length and tiller number, no significant difference was observed with *krp3-1*, unlike Tp309, which had a significantly longer internode (19%) as well as higher tiller number (38%) (**Fig. 4C, D**). Like *krp3* mutants, mutation of *mpk3* or *krp3mpk3* also did not affect leaf size (**Sup. Fig. 10E, F, Fig. 4E**). Minor reduction in maximum seed per panicle was also observed in *mpk3-1* and *mpk3-2* mutant lines (8%) compared to Tp309 (**Sup. Fig. 10G**). The seed number reduction was more pronounced in *krp3mpk3* line (11%) compared to Tp309, which is similar to *krp3-1* (**Fig. 4F, G**). While seed number got reduced, no change in seed weight was detected either in *mpk3* mutants or *krp3mpk3* double mutants compared to Tp309 (**Sup. Fig. 10H**, **Fig. 4H)**. Overall, the observations suggested that KRP3 and MPK3 function in a linear pathway in regulating plant growth and development.

**Figure 4:**
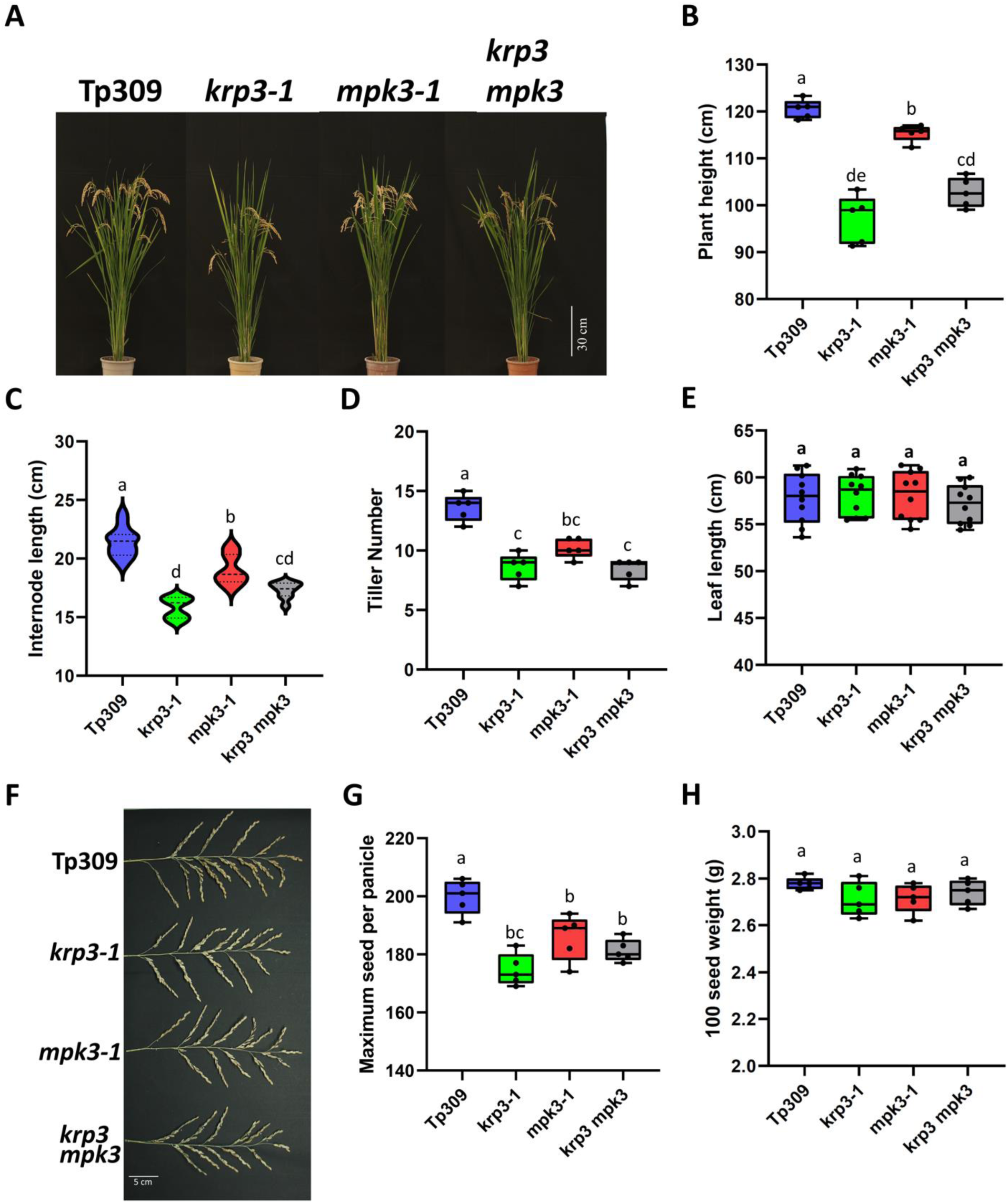
KRP3 and MPK3 work in compliance. **A.** Pictorial representation of Tp309, *krp3-1*, *mpk3-1*, *krp3mpk3* mature plants respectively. **B-D.** Box plot and violin plot representing the plant height (B) (n=5), tiller number per plant (C) (n=5) and 1^st^ internode length (D) (n>20) of all the lines. In all three parameters *krp3-1*, *mpk3-1* and *krp3mpk3* showed a similar trend of reduced plant height, internode length as well as tiller number. **E.** Mature leaf length of all the lines. The leaf length of all three mutant lines was comparable to Tp309. **F.** Comparative image of a mature panicle of all the lines. **G.** Maximum seed number of the primary tiller of all the lines. All three mutant lines showed lower seed numbers in the primary panicle. **H.** Box plot showing the 100 seed weight of Tp309, *krp3-1, mpk3-1* and krp3mpk3 lines indicating no significant difference. One-way ANOVA and Tukey’s multiple comparisons test was conducted to compare the differences. Different alphabets represent significant changes observed in Tukey’s multiple comparisons test.

### MPK3 assists KRP3 in maintaining cell division rate

To elucidate the effect of *mpk3* and *krp3mpk3* knockout conditions in rice seedling growth, shoot and root lengths of fourteen-day-old seedlings were evaluated. It was found that the *mpk3* mutant has minimal effect on shoot and root length reduction (**Sup. Fig. 11A-C**). When shoot and root length of *krp3mpk3* seedling were measured and compared with Tp309, *krp3-1* and *mpk3-1,* it was evident that *krp3mpk3* had a significant reduction in both shoot (14%) and root length (29%) (**Fig. 5A-C**). It was further observed that *krp3-1* and *krp3mpk3* shoot and root length were statistically indifferent (**Fig. 5A-C**). The root cell elongation pattern of *krp3-1* and *krp3mpk3* were similar to each other, showing 1.6-fold longer cell length than that of Tp309 (**Fig. 5D**). *mpk3-1* also exhibited moderately longer cell length (1.2-fold) to that of Tp309 (**Fig. 5D**). Though all mutants showed higher cell length, cell width difference was not significant (**Fig. 5E**). Rate of cell division using EdU labelling in *mpk3-1* and *mpk3-2* root tips was checked and considerably a smaller number of cells were found to be undergoing S phase compared to Tp309 (**Sup. Fig. 11D, E**). Similarly, cells undergoing the S phase in *krp3mpk3* were checked, and 30% lower EdU-labeled cells were observed as compared to Tp309 (**Fig. 5F, G**). No change in cell number undergoing S phase was observed between *krp3-1* and *krp3mpk3* double mutant (**Fig. 5F, G**). The effect of HU in root elongation was compared among Tp309, *krp3-1*, *mpk3-1* and *krp3mpk3*. Root length reduction rate showed *krp3-1* and *krp3mpk3* had similar hypersensitive effects towards S phase inhibition compared to Tp309, while *mpk3-1* exhibited a completely contrasting phenotype of low sensitivity to HU (**Fig. 5H**). The subsequent effect of HU on cell elongation of mutant lines was measured, and an antagonistic trend to that of root length reduction was found. *mpk3-1* showed the highest elongation of cells while *krp3-1* and *krp3mpk3* showed the lowest root cell elongation compared to Tp309 (**Fig. 5I**). The findings suggested that MPK3 helps KRP3 to maintain the rate of cell division.

**Figure 5:**
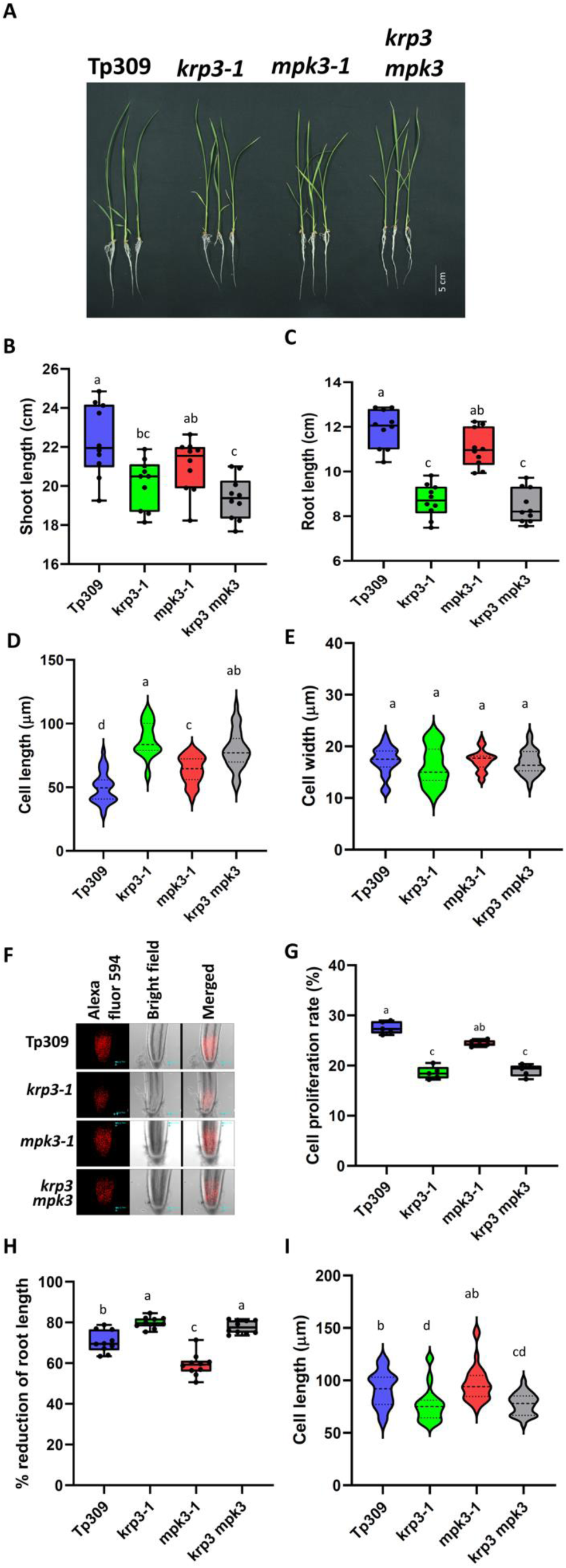
KRP3 and MPK3 function in a linear order to maintain cell division rate. **A.** Image showing fourteen-day-old seedling of Tp309, *krp3-1*, *mpk3-1*, *krp3mpk3*. **B, C.** Shoot (B) and root length (C) of fourteen-day-old seedlings of all the lines. A strong reduction in root and shoot length was observed in *krp3-1* and *krp3mpk3* lines. *mpk3-1* also showed a similar trend but with a moderate effect. **D, E** Violin plot showing mature root cell length (D) and width (E) of all the lines. Cell width was similar in all four lines while *krp3-1 mpk3-1* and *krp3mpk3* showed longer cell length compared to Tp309. **F.** Image of EdU-Alexa fluor 594 labeled root tip of Tp309, *krp3-1*, *mpk3-1* and *krp3mpk3* observed under confocal microscope. **G**. Quantitative box plot showing the rate of cell proliferation at root division zone. *krp3-1* and *krp3mpk3* showed similar reductions in cell proliferation compared to Tp309. **H.** Percent root length reduction in the presence of HU in the seedlings of all the lines. *mpk3-1* showed less sensitivity towards HU while *krp3mpk3* showed hypersensitivity like *krp3-1* compared to Tp309. **I.** Mature cell length of seedling root grown in the presence of HU. A similar but opposite trend to that of root length reduction was observed where *mpk3-1* showed longer cell length but *krp3-1* and *krp3mpk3* showed reduced cell length with respect to Tp309 cell length. One-way ANOVA and Tukey’s multiple comparisons test was conducted to compare the differences. Different alphabets represent significant changes observed in Tukey’s multiple comparisons test.

### Phosphorylation of KRP3 leads to sturdier cell cycle inhibition

To evaluate the effect of KRP3 phosphorylation on plant growth and yield, KRP3 phospho-dead (KRP3^S17A, S82A^) and phospho-mimetic (KRP3^S17E, S82E^) overexpressing transgenic lines were developed. Three independent lines, showing comparable KRP3 overexpression with KRP3-OE lines, for both KRP3^S17A,S82A^ and KRP3^S17E,S82E^, and named as *KRP3^AA^-OE1/2/3* and *KRP3^EE^-OE1/2/3*, respectively (**Sup. Fig. 12A, B**). Initially, mature plant height and internode length for both KRP3^AA^-OE and KRP3^EE^-OE lines were checked. Significant reduction in plant height (13-6% and ∼22%) and internode length (7-12% and ∼21%) was observed in KRP3^AA^-OE lines as well as KRP3^EE^-OE lines compared to Tp309 (**Sup. Fig. 13A-F**). When plant height of *KRP3-OE1*, *KRP3^AA^-OE1* and *KRP3^EE^-OE1* was compared with Tp309 and VC, it was clear that *KRP3^EE^-OE1* exhibited the highest reduction in plant height (21%) followed by *KRP3-OE1* (15%) and *KRP3^AA^-OE1* (13%) (**Fig. 6A, B**). When the reduction in internode length was compared among these lines, *KRP3^AA^-OE1* showed the least reduction in internode length (12%) while *KRP3^EE^-OE1* exhibited the highest reduction (25%) while *KRP3-OE1* showed an intermediate effect (20%) (**Fig. 6C**). A contrasting trend was observed in tiller number of KRP3^EE^-OE overexpression lines. KRP3^EE^-OE lines showed a significantly higher (50%) number of tillers compared to Tp309 and VC plants, while KRP3^AA^-OE lines showed no significant changes (**Sup. Fig. 13G, H**). When compared among the three forms of overexpression lines, it was observed that KRP3-OE1 or KRP3^AA^-OE1 had no significant effect on tiller number while KRP3^EE^-OE1 positively influenced tiller formation (**Fig. 6D**). Leaf length also showed a similar trend like plant height, KRP3^AA^-OE lines, as well as KRP3^EE^-OE lines, showed significant reduction (∼10% and 25-18% respectively) in leaf length (**Sup. Fig. 13I, J**). Comparing each other it was clearly visible that *KRP3^EE^-OE1* had the shortest leaf followed by *KRP3-OE1* and *KRP3^AA^-OE1*, while leaf width was unaffected (**Fig. 6E-G**). KRP3^AA^-OE lines showed inconsistency in the highest seed number per panicle. *KRP3^AA^-OE1* and *KRP3^AA^-OE2* showed significant reduction (19% and 9%) in seed number while no significant change in *KRP3^AA^-OE3* was observed (**Sup. Fig. 14A**). On the other hand, all the lines studied for KRP3^EE^-OE showed a significant reduction in maximum seed number per panicle (41-37%) (**Sup. Fig. 14B**). *KRP3^EE^-OE1* showed the highest reduction in seed number (41%) while *KRP3^AA^-OE1* had least reduction (19%) (**Fig. 6H, I**). Lower seed weight was observed in all KRP3^AA^-OE, and KRP3^EE^-OE lines (**Sup. Fig. 14C, D**). Reduction in mature seed length was observed in KRP3^AA^-OE, and KRP3^EE^-OE lines, while seed width was only reduced in KRP3^EE^-OE lines (**Sup. Fig. 14E - H**). *KRP3^EE^-OE1* showed extreme reduction in seed weight (21%), length (11%) and width (6%), flowed by *KRP3-OE1* (8%, 7% and 1%) and *KRP3^AA^-OE* (4%, 5% and 0.6%) (**Fig. 6J-M**).

**Figure 6:**
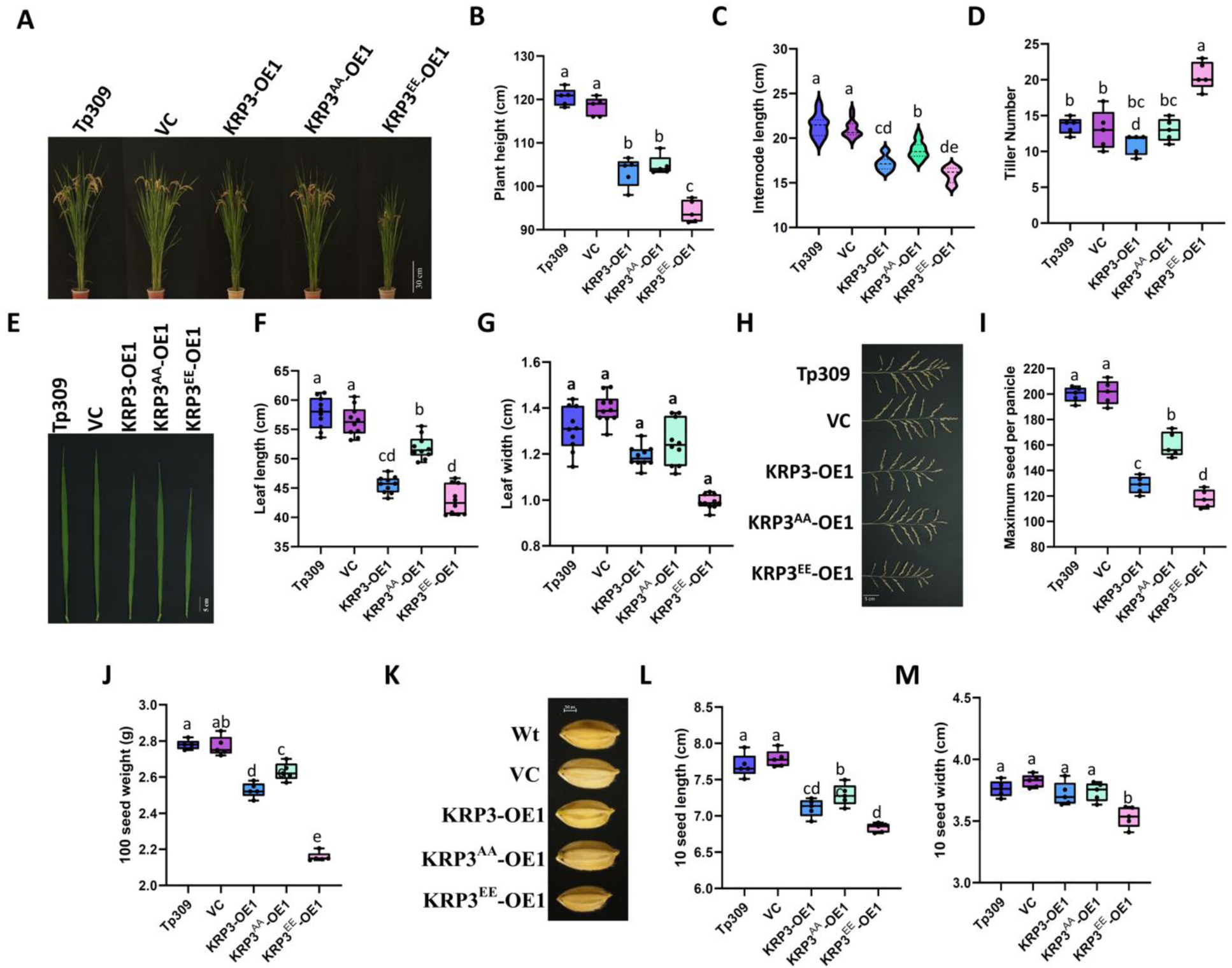
Overexpression of KRP3^EE^ promotes stronger plant growth inhibition. **A.** Pictorial representation of Tp309, VC, KRP3-OE1, KRP3^AA^-OE1 and KRP3^EE^-OE1 mature plant. **B, C.** Box and violin plot representing the plant height (B), and 1^st^ internode length (C) of mature plants of all the lines. Reduction in plant height and tiller number was observed in all three types of OE lines. KRP3^EE^-OE showed the highest reduction and KRP3^AA^-OE showed the least reduction in plant height and internode length. **D.** Tiller number of all the lines (n=5). Though the tiller number in KRP3-OE1 and KRP3^AA^-OE1 was unaltered, the KRP3^EE^-OE1 line showed a higher number of tillers per plant. **E.** Photograph representing fully grown last vegetative leaf of all the lines. **F, G.** Leaf length (F) and width (G) of all the lines. Like KRP3-OE1, KRP3^AA^-OE1 and KRP3^EE^-OE1 showed a reduction in leaf length and no difference in leaf width. **H.** Comparative photograph of a mature panicle of all the lines. **I, J.** Maximum seed number of the primary tiller (I) and 100 seed weight (J) of all the lines. KRP3-OE1, KRP3^AA^-OE1 and KRP3^EE^-OE1 showed a reduction in seed number and seed weight among which KRP3^EE^-OE1 showed the strongest reduction in seed number and seed weight. **K.** Pictorial representation of seed size of Tp309, VC, KRP3-OE1, KRP3^AA^-OE1 and KRP3^EE^-OE1. **L, M.** Box plot showing seed length (L) and width (M). Justifying seed weight KRP3^EE^-OE1 showed a reduction in both seed length as well as width, while KRP3-OE1 and KRP3^AA^-OE1 only showed seed length reduction. One-way ANOVA and Tukey’s multiple comparisons test was conducted to compare the differences. Different alphabets represent significant changes observed in Tukey’s multiple comparisons test.

Comparing shootand root lengths of seedlings, a significant reduction in both shoot and root lengths was observed in KRP3^AA^-OE lines (19-29% and 20-29%) as well as KRP3^EE^-OE lines (23-37% and 39-49%) (**Sup. Fig. 15A-F**). In the case of seedling shoot length, *KRP3-OE1*, *KRP3^AA^-OE1* and *KRP3^EE^-OE1* were not different among themselves, while they were significantly shorter than Tp309 and VC seedlings (33%, 29% and 37%) (**Fig. 7A, B**). Comparing root length, it was observed that *KRP3-OE1*, *KRP3^AA^-OE1* and *KRP3^EE^-OE1* roots are not only shorter than Tp309 and VC seedlings, but there is also significant difference among each other. It was evident that *KRP3^EE^-OE1* root is the shortest, followed by *KRP3-OE1* and *KRP3^AA^-OE1* (**Fig. 7C**). *KRP3^EE^-OE1* also exhibited the longest and widest root cell while *KRP3^AA^-OE1* had no significant effect in root cell elongation compared to Tp309 and VC (**Fig. 7D, E**). Cell division rate measured using EdU incorporation assay depicted that *KRP3^EE^-OE1* has the strongest inhibition in cell division (66% less cell division) followed by *KRP3-OE1* (48%), while *KRP3^AA^-OE1* had minor effect (24%) on cell cycle inhibition (**Fig. 7F)**. *KRP3^AA^-OE1*, showed moderate hypersensitivity to HU, while *KRP3-OE1* and *KRP3^EE^-OE1* showed less sensitivity compared to Tp309 and VC (**Fig. 7G**). Cell elongation in the presence of HU was comparable within Tp309, VC, *KRP3-OE1*, *KRP3^AA^-OE1* and *KRP3^EE^-OE1*, where only KRP3^EE^-OE1 showed moderately higher cell length compared to others (**Fig. 7H, I**). The data revealed that the phosphorylation of KRP3 by MPK3 leads to substantial inhibition of the cell cycle.

**Figure 7:**
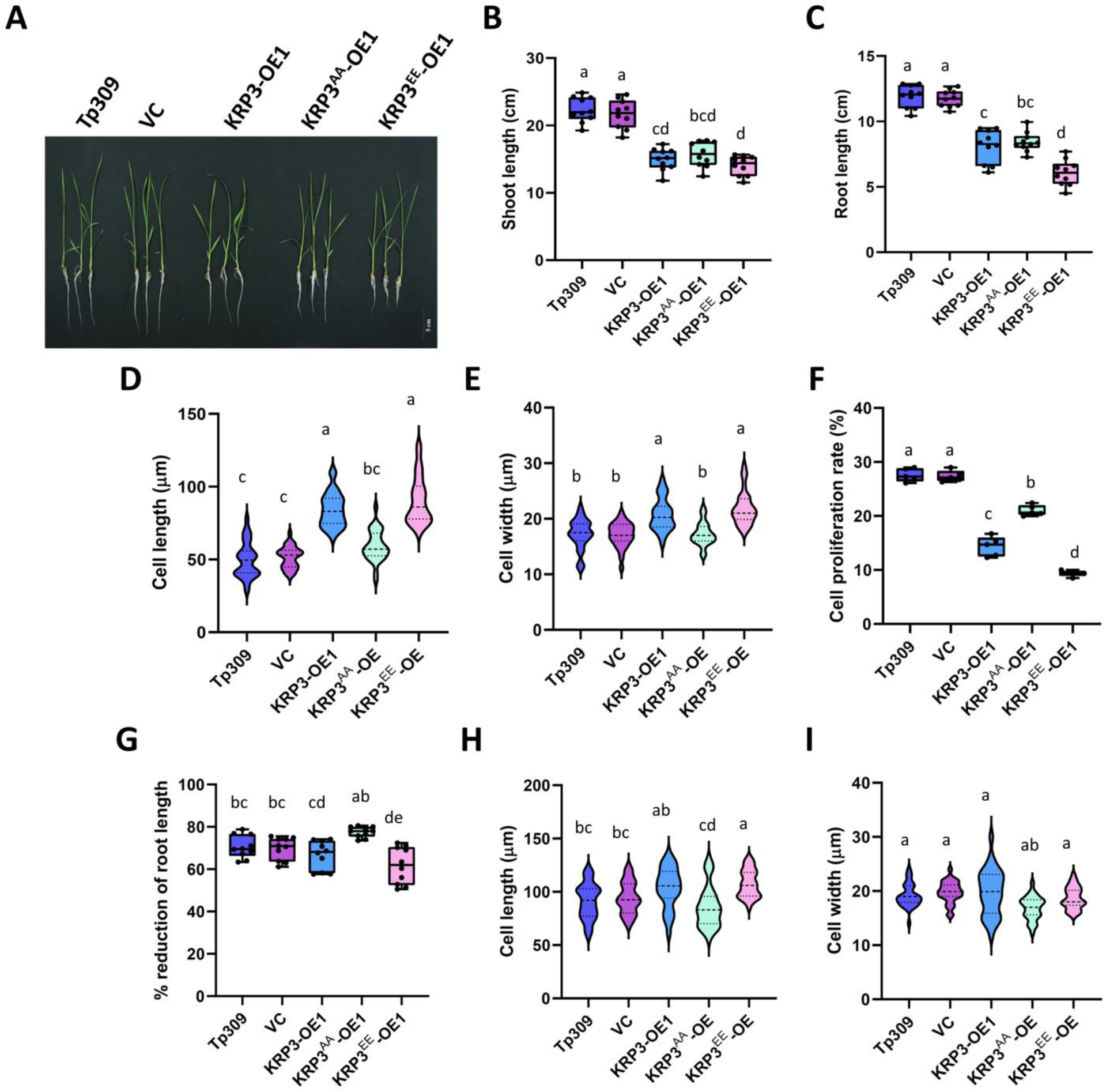
KRP3 maintain cell division rate by regulating G-S checkpoint. **A.** Image depicting fourteen-day old seedling of Tp309, VC, KRP3-OE1, KRP3^AA^-OE1 and KRP3^EE^-OE1. **B, C.** Shoot (B) and root length (C) of fourteen days old. All representative overexpression lines showed a reduction in shoot and root length. **D, E** Mature root cell length (D) and width (E) of all the lines. KRP3^EE^-OE1 exhibited the longest and widest cells followed by KRP3-OE1 and KRP3^AA^-OE1 compared to Tp309 and VC. **F**. Rate of cell division at root division zone. KRP3^EE^-OE1 showed the strongest inhibition of cell proliferation. **G-I.** Percent root length reduction (G), cell length (H) and width (I) in the presence of HU in the seedlings of all the lines. KRP3^AA^-OE1 showed moderate sensitivity while KRP3^EE^-OE1 showed moderate tolerance. One-way ANOVA and Tukey’s multiple comparisons test was conducted to compare the differences. Different alphabets represent significant changes observed in Tukey’s multiple comparisons test.

### Phosphorylation of KRP3 at S17 and S82 position reduces its turnover rate

To identify the exact mechanism how phosphorylation is affecting KRP3’s function, the localization of KRP3, KRP3^AA^ and KRP3^EE^ was checked in *N. benthamiana* leaf. mCherry tag was added in the amino-terminal side of KRP3 and its mutant forms. NLS-tagged mGFP was used as a nuclear marker. While mCherry alone was found in both nucleus as well as cytosol, KRP3 and its mutant forms were found to localize specifically in the nucleus (**Sup. Fig. 16**). Effect of phosphorylation on KRP3’s interaction with CDKA or CycD was also checked. BiFC interaction assay was conducted where KRP3, KRP3^AA^ and KRP3^EE^ cloned with the C-terminal domain of YFP was co-infiltrated with CDKA or CycD cloned with the N-terminal of YFP. All the constructs were also infiltrated along with complimenting empty vector controls. No effect on KRP3’s interaction with CDKA or CycD was observed (**Sup. Fig. 17**). The effect of phosphorylation on KRP3 protein stability was also checked using cell-free degradation assay. MBP-KRP3-6xHis, MBP-KRP3^AA^-6xHis and MBP-KRP3^EE^-6xHis chimeric proteins were incubated for 120 min at 4°C with crude plant protein extract of Tp309 seedling. Western blot analysis using anti His-antibody showed that the content of KRP3, as well as KRP3^AA^ protein, were reduced drastically while the reduction of KRP3^EE^ protein level was significantly less (**Fig. 8A**). The observation validated that the reason for stronger inhibition of cell cycle in the transgenic line over-expressing phospho-mimetic version of KRP3 is the stabilization of KRP3 protein turnover post phosphorylation by MPK3.

**Figure 8:**
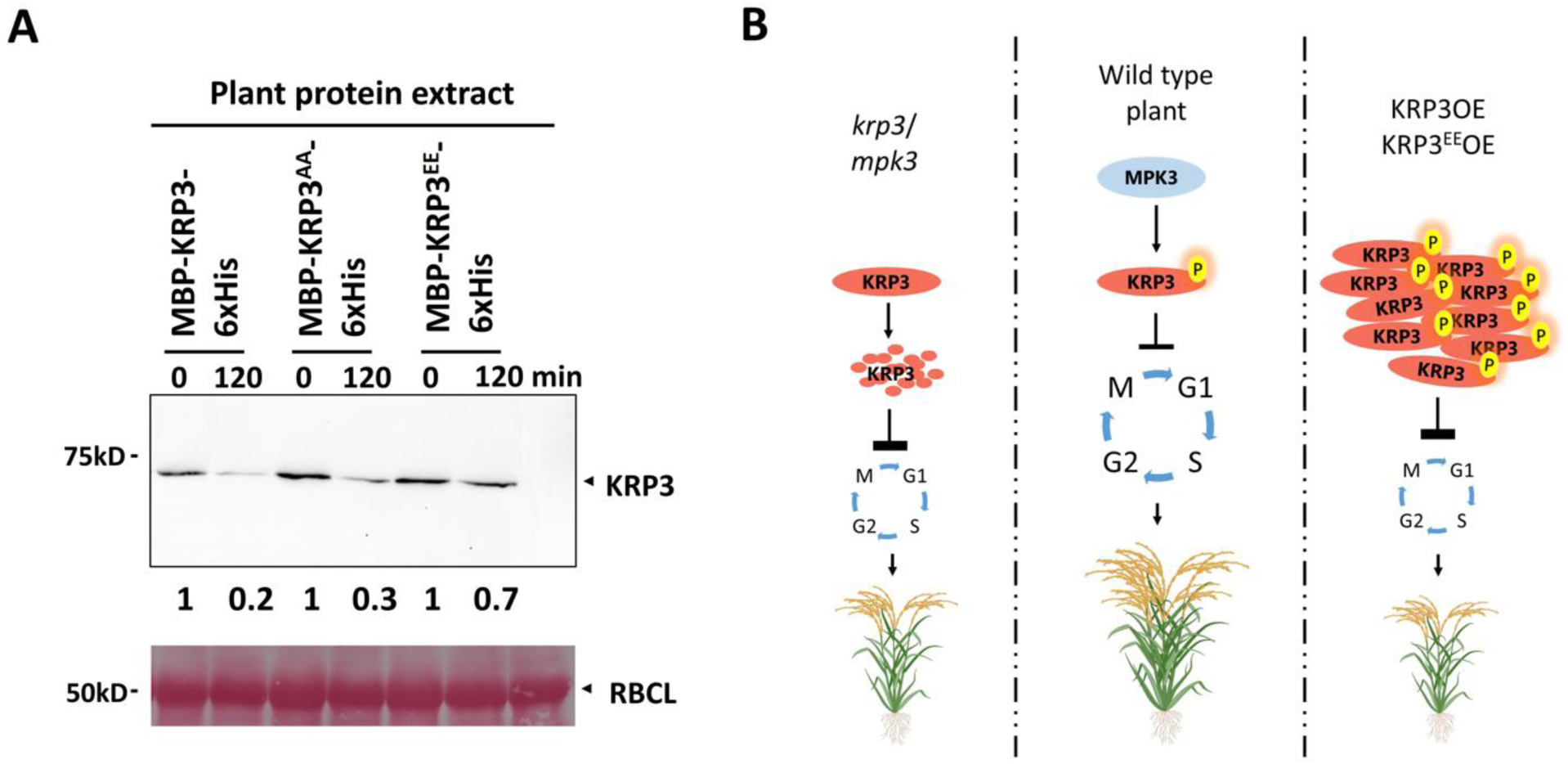
Phosphorylation of KRP3 at S17 and S82 positions promotes its stability. **A.** Cell-free degradation assay of KRP3, KRP3^AA^ and KRP3^EE^, incubated with plant protein extract for 120 min. Western blot showing KRP3 protein level of initial and post 120 min of incubation. KRP3^EE^ was found to be the most stable form compared to KRP3 and KRP3^AA^ **B**. Model for KRP3 function, maintaining plant vigor and yield in a dose and phosphorylation-dependent manner.

## Discussion

Cell division and subsequent elongation are integral to plant growth and development. Studies with cell cycle inhibitors in plants have shown their role in regulating plant morphogenesis. Ectopic overexpression of rice *KRP1* has shown increased cell size and reduced cell division, leading to smaller leaves and ill-developed seeds with altered size and ploidy levels in the embryo (Ajadi et al., 2020; Ren et al., 2008). Other than *KRP1*, *OsiICK6* or *KRP4* overexpression in *indica* rice variety strongly reduced organ size and grain production (Xu et al., 2023; Yang et al., 2011). Although in *Arabidopsis*, single or double mutation of *KRP* genes has a negligible effect on overall plant morphology, in rice, the story is quite different (Cao et al., 2018). Mutation of KRPs in rice like *krp1* or *krp5* has shown a significant effect on seed filling rate (Ajadi et al., 2020; Xu et al., 2023). Recent studies indicate the difference in the role of KRPs between dicot and monocot plants. Phylogenetic analysis of KRPs from monocot genera yielded some critical observations. Interestingly, *KRP3* homologs in monocots have only been identified in the Poaceae family. This family comprises the most economically important cereal crops like rice, maize, wheat, etc. Looking at the importance of rice, the probable role of *KRP3* in maintaining rice morphogenesis and productivity was studied. Phenotypic analysis of two independent *krp3* knock-out mutants showed reduced plant height and low tiller formation. Meticulous measurement of internode length revealed that reduction in plant height is a consequence of shorter internodes. Reduction in plant height and tiller number of crop species is considered as one of the superior traits in modern agriculture, where overall seed production per tiller is increased (Liu et al., 2018; Takai, 2023). KRP3 was also observed to play an important role in maintaining crop yield*. krp3* mutation negatively affected the seed number per panicle. Interestingly, in *krp3-1* and *krp3-2,* though seed number was reduced , seed size and weight were unaffected. Traits such as effective tiller number, seed number, seed size and weight are some of the key components determining rice yield (Li et al., 2021). Thus, with a reduced tiller and seed number, *krp3* plants exhibited a significant reduction in rice productivity. These observations hint towards the role of KRP3 in the regulation of crop architecture and yield.

*KRP3* overexpression lines also showed an overlapping phenotype with that of *krp3* mutants. All three studied overexpression lines of *KRP3* showed a negative effect on plant height, leaf length, tiller number, seed number and seed weight. Displaying a similar effect in both overexpression as well as in mutant of any gene is not unlikely and has been observed in plenty of cases where a gene executes its function in a dose-dependent manner (Chen et al., 2023; Spadafora et al., 2012). Considering KRP3 as a negative regulator of the cell division cycle, justifies the reduction in overall plant size and productivity in KRP3 overexpression lines. Similar growth compromise has been observed in both *Arabidopsis* and rice in case of overexpression of KRP genes (Barrôco et al., 2006). However, the cause of growth and yield penalty in *krp3* plants were not clear.

While KRP3 influences rice architecture, it was important to understand whether it was through modulating cell division rate or mature cell size. EdU is an analogue of thymidine, and it can be utilized to detect cell proliferation rates by externally supplying it in the growth media (Chehrehasa et al., 2009). The estimated cell division rate in the growing root tips of *krp3-1* and *krp3-2* revealed that mutation of *krp3* negatively affects cell cycle progression. This reduction in cell division led to cell elongation in *krp3-1* and *krp3-2* compared to Tp309. This observation is consistent with earlier studies reporting that the inhibition of cell division leads to extended cell elongation (Song et al., 2022). *krp3* mutants also exhibited reduced root length, hinting that the elongation of root cell is not sufficient to surpass the effect of lower cell division in the dividing zone. Interestingly, the rate of cell cycle inhibition and cell elongation in *krp3* is similar to that of KRP3 overexpression lines.

Quantitative analysis of *KRP3* transcript abundance using the cell cycle inhibitors showed that KRP3 only participates in S phase checkpoint regulation. Hydroxyurea (HU) and Mimosine, two well-known inhibitors of ribonucleotide reductase, were used as S-phase inhibitors (Pan et al., 2021). Inhibition of ribonucleotide reductase leads to the depletion of deoxyribonucleic acid, which is crucial for DNA replication, resulting in the activation of the S phase checkpoint and inhibition of the cell cycle (Musiałek and Rybaczek, 2021). For M phase inhibition, Propyzamide and Oryzalin, two anti-mitotic drugs inhibiting microtubule polymerization, were used (Nakamura et al., 2004). The fact that KRP3 is involved in S phase checkpoint regulation was also evident through hypersensitivity towards HU. Similar inhibition in root growth in the presence of HU has also been reported in mutants of other S phase checkpoint regulators like *atm*, *atr*, and *wee1* (De Schutter et al., 2007; Pedroza-Garcia et al., 2021). Though *krp3* mutants were hypersensitive to HU, root length reduction in KRP3 overexpression lines was identical to Tp309, indicating a rate limitation in KRP3 function, possibly pointing towards a post-translational modification.

Post-translational modification like phosphorylation has been known for a long to maintain the functionality of proteins (Verma et al., 2020). Recent studies have demonstrated that three MAP Kinases, MPK3, MPK4 and MPK6, phosphorylate the E2F2 transcription factor and inhibit its function in the presence of HU (Singh et al., 2023). Out of these three MPK Kinases, MPK3 specifically interacted with KRP3 in the nucleus in our study. The interaction of proteins with MPK3 often leads to its phosphorylation (Bhagat et al., 2022a). We observed that MPK3 activated by AtMKK4DD can phosphorylate KRP3 and MBP. It was also observed that bacterially expressed MPK3 showed auto and trans-phosphorylation, as MPK3, without activation by upstream MAPKK, can phosphorylate MBP as well as KRP3. Auto-activation properties of bacterially expressed MAP Kinases have been reported previously (Sethi et al., 2014; Singh and Sinha, 2016). Previous reports also suggest that MAP Kinase phosphorylates its substrate at the S/TP motif (Dóczi and Bögre, 2018). Four SP sites were identified as possible MAP Kinase phosphorylation sites in the KRP3 protein sequence. These putative phosphorylation sites were validated by using five combinations of KRP3^S>A^ constructs as Ala is considered a non-phosphorylatable amino acid (Dóczi and Bögre, 2018). *In-vitro* kinase assay using these mutant forms suggested that MPK3 phosphorylates KRP3 at Ser 17 and S82 positions. This was further validated using a KRP3^S17A, S82A^ (KRP3^AA^) mutant and this mutated form of KRP3 also did not get phosphorylated by MPK3.

MPK3 regulate multiple cell cycle checkpoint proteins like WEE1 and SMR1 (Banerjee et al., 2023). Thus, MPK3-mediated phosphorylation of KRP3 must be an essential regulation point in the rice cell cycle. The present study evaluated the role of MPK3 in rice morphogenesis and yield using *mpk3* knock-out mutants. Both *mpk3-1* and *mpk3-2* behaved similarly to the *krp3* mutant phenotype, with reduced plant height, shorter internode length, and a smaller number of tiller and seed in the primary panicle. Though root growth of *mpk3* mutants showed no difference in normal conditions, it showed less sensitivity to HU-mediated cell division inhibition. In the absence of MPK3, the checkpoint regulation at the G1-S phase might be hampered as it is also involved in the regulation of other heck point regulators (Banerjee et al., 2023). The *krp3mpk3* double mutant also exhibited a similar trend in vegetative growth parameters like reduction in plant height, internode length and tiller number. Interestingly, there was no significant difference between *krp3-1* and *krp3mpk3* mutant lines. Similar trends were observed in the case of seed number reduction and shoot and root length reduction in fourteen-day seedlings. *krp3mpk3* also showed hyper HU sensitivity and reduced cell division rate like *krp3-1*. This similar trend of mutant phenotype of *krp3-1*, *mpk3-1* and *krp3mpk3* indicates that KRP3 and MPK3 are involved in a single pathway regulating rice morphogenesis and yield. These observations further strengthened the role of MAPK-mediated phosphorylation of KRP3 in cell cycle regulation.

Phenotyping of constitutive overexpression transgenic lines of KRP3 phospho-dead (KRP3^AA^-OE) and phospho-mimetic (KRP3^EE^-OE) form provided insights into the role of its phosphorylation in plant architecture and yield regulation. For phospho-mimetic form of KRP3, Ser was converted to Glu (KRP3^EE^), which mimics the negative charge imposed due to phosphorylation (Dóczi and Bögre, 2018). A clear difference between KRP3^EE^ and KRP3^AA^ overexpression lines was observed, where KRP3^EE^-OE lines exhibited a stronger reduction in plant height, internode length and leaf length. This phenotype indicates that phosphorylation of KRP3 increases its potentiality to inhibit the cell cycle. Altogether, it can be concluded that KRP3 play a crucial role in maintaining rice vigor and yield. The phosphorylation of KRP3 is critical to regulating the KRP3 protein turnover rate. It is also evident that the presence of KRP3 in a checked volume in dividing cells is the decisive factor for cell division rate.

MAP Kinase signaling had diverse effects on its substrates due to phosphorylation, which includes modulating localization, gain or loss of function, or even affecting stability (Bhagat et al., 2022b; Kim et al., 2022). Some KRP proteins contain an NLS signal and get localized into the nucleus (Boruc et al., 2010). Rice KRP3 was also found to be localized in the nucleus. However, similar to KRP3, its phospho-dead, and phospho-mimetic versions were also found to be localized into the nucleus, indicating phosphorylation doesn’t regulate the localization of KRP3. Further, phosphorylation doesn’t seem to regulate the interaction of KRP3 with its target proteins. It is known that KRP3 interacts with the CDKA-CycD complex and inhibits its function (Mizutani et al., 2010). No change in the interaction between KRP3, KRP3^AA^ and KRP3^EE^ forms with CDKA and CycD was observed. Another aspect that regulates the function of KRPs is the protein turnover rate. Multiple mechanisms involving proteasomal and non-proteasomal mediated degradation governs the protein-level maintenance of KRPs (Kim et al., 2008; Li et al., 2016; Noir et al., 2015). MAP kinase-mediated phosphorylation also has implications for regulating protein turnover (Kim et al., 2022). In the case of KRP3, the cell-free degradation assay clearly showed that the stability of KRP3^EE^ is significantly enhanced over KRP3 and KRP3^AA^, which indicates that phosphorylation of KRP3 at S18 and S72 positions protects it from degradation.

Adequate protein level maintenance of some KRPs, in a dividing cell, is crucial in *Arabidopsis*. In the case of AtKRP4, it has been observed that the parental inherited level of KRP4 protein regulates the initiation and time duration of subsequent cell division (D’Ario et al., 2021). Other than *AtKRP4,* overexpression of *AtKRP6* has been shown to promote cell division in roots infected by root-knot nematode (Vieira et al., 2014). With this information, we relooked at our findings to validate that KRP3 regulate rice cell division in a dose-dependent manner. From the plant vegetative growth and yield parameters, it is evident that *krp3* mutant and *KRP3-OE* or *KRP3^EE^-OE* are showing similar phenotypes in several traits like plant height, root length, seed number, cell length, cell division rate, etc. **Fig 8B** represents a simplified model depicting the effect of KRP3 phosphorylation by MPK3 and its impact on rice plants. In conditions where KRP3 protein cannot get phosphorylated, i.e. *mpk3* or KRP3^AA^-OE, similar deviation from normal phenotype was observed, but the reduction in plant growth and yield was less severe. This demonstrates that KRP3 functions in a dose-dependent manner to maintain cell division rate and cell size in rice-growing organs and requires MPK3-mediated phosphorylation for its stability. Phosphorylated KRP3 modulates cell proliferation in rice to maintain its plant architecture and yield contributing traits.

## Material and Methods

### Plant Growth Conditions

Japonica rice (*Oryza sativa* Lin.) variety Taipei309 (Tp309) has been used in this study. Surface sterilized seeds were germinated in the dark and post-germination seeds were transferred in 1/4^th^ strength MS media (pH 5.8), and kept for growing under 16h light 8h dark at 28° C for fourteen days and used for shoot and root length measurement. For estimating yield-related parameters plants were grown in greenhouse with a 16h light and 8h dark photoperiod cycle at 28 ± 2° C temperature with 60% relative humidity. 10” planter filled with nutrient-reach soil were used for growing plants.

### Development of phylogenetic tree

Sequences of one hundred and forty-four monocot specific KRP proteins were retrieved from NCBI database. The phylogenetic tree was constructed using Maximum Likelihood method and JTT matrix-based model, with 100 bootstrap replicates. MEGA11 platform was used for developing the tree (Tamura et al., 2021).

### Generation of knockout rice mutant, transgene overexpressing plants and promoter-reporter lines

For the generation of *krp3* and *mpk3* knockout mutants using the CRISPR-Cas9 tool, single guide RNA was designed using CRISPR-P, and cloned into pRGEB32 vector. For *krp3mpk3* double knockout mutant gRNA was cloned in tandem with tRNA (as described in Xie et al., 2015) in pRGEB32 vector. The constructs were transformed into *A. tumefaciens* strain EHA103 to develop transgenic lines. Transgenics were raised as previously described (Rengasamy et al., 2024) Homozygous mutation was detected by DNA sequencing using Sanger method. Lines with homozygous mutation and transgene (Cas9 and HPT) free in T1 generation were selected for next-generation propagation.

For the development of over-expression lines, KRP3, KRP3^AA^ and KRP3^EE^ fused with N-term 3xFLAG and C-term 4xMyc tag, were cloned under CamMV-35S promoter and NOS terminator, in pCAMBIA1300 vector. The developed construct and empty pCAMBIA1300 vector were transformed in EHA103, followed by stable transgenic development.

For developing promoter-reporter lines, 2kb region of KRP3 gene upstream to the coding sequence was amplified and cloned into pCAMBIA1301 using EcoRI and NcoI restriction sites, replacing CamMV-35S promoter. The developed construct and pCAMBIA1301 vector devoid of CamMV-35S promoter were transformed in EHA103, followed by the development of stable transgenic lines. Primers used for the development of constructs are mentioned in supplementary table 1.

### Measurement of plant height and tiller number

Plants were grown in greenhouse condition till maturity. After maturity, five plants from each line were uprooted, soil was washed, and photographed. Height was measured from the root initiation zone to the panicle end using ImageJ software. The primary tiller numbers of five mature plants were counted manually.

### Internode length, leaf length and width measurement

Internodes were exposed by removing all leaves form the primary tiller and then photographs were taken of all tillers from five mature plants. 1^st^ internode (identified as mentioned in Ji et al., (2019)) length of twenty internodes was measured using ImageJ software. Similarly, fully grown last vegetative leaf (leaf immerged prior to flag leaf) length was measured (from leaf tip to ligule) using ImageJ software.

### Measurement of maximum seed per panicle, seed weight and seed size

Seed numbers in the highest seed-bearing panicle of five plants were counted manually. For seed weight, a hundred seed weights were counted and weighed using a weighing machine (Sartoris, TE64). Seed length and width were measured using ImageJ, by taking photographs of ten seeds arranged in a line. Length and width were recorded from five independent plants of every line.

### Estimation of seedling shoot and root length

Fourteen-day old seedlings, grown in 1/4^th^ strength MS media (pH 5.8) under 16h light and 8h dark conditions at 28°C, were photographed. Shoot and root length of twenty seedlings were measured for every line using ImageJ software.

### Estimation of cell length and width

Around 1 cm long root portion from the mature zone (below crown region) of seven-day-old seedlings was cut and incubated with 1 μg/mL Propidium Iodide (PI) for 20 min. Post incubation samples were washed three times with water and visualized under a confocal microscope (SP8, Leica) for PI fluorescence. Cell wall autofluorescence was also captured as described by Lahlali et al., (2016) for better evaluation. Cell length and width were measured using Leica LAS AF Lite software.

### GUS staining

Seven day old Seedlings and 1st internode region of ProKRP3::GUS and VC-GUS lines were incubated in GUS staining solution (200µM of X-Gluc, 0.2mM K_3_Fe(CN)_6_, 0.2mM K_4_Fe(CN)_6_, 100mM EDTA, 0.1% TritonX100, pH-7.0) for 12h. After staining, samples were washed for three times in a de-staining solution containing ethanol, acetic acid and glycerol (in a ratio of 3:1:1). After washing, photographs were taken using a Canon DSLR camera or Stereo zoom microscope with a camera (Zeiss).

### EdU incorporation assay

To visualize the rate of cell proliferation EdU (5-Ethynyl-2-deoxyuridine) incorporation assay was used. EdU was added in liquid growth media with a final contrition of 50µM for incorporation into the growing root tip. Root tips were harvested after 4h of EdU addition and fixed into a fixative solution (4% paraformaldehyde, 0.1% TritonX-100, in PBS pH 7.4) for 1h. After fixation, samples were washed with PBS for three times and then incorporated EdU was labelled with Alexa Fluor 594 dye using Click-iT™ EdU Cell Proliferation Kit (Invitrogen) as per manufacturer’s protocol. Root tips were further washed using PBS three times and used for confocal microscopic imaging (SP8, Leica). The fluorescence of Alexa Fluor 594 was detected with an excitation wavelength around 590 nm and an emission wavelength around 618 nm.

### Determination of cell proliferation rate

For determination of cell proliferation rate, roots of seven-day-old seedlings were incubated with 50µM EdU for 1h followed by fixation and labelling. Post labelling, root tips were incubated with a cell maceration cocktail (1% cellulose, 1% pectinase in PBS pH 7.4) for 1h. Later, root tip cells were squashed on a slide and counter stained with DAPI for nuclear staining. Slides were visualized under confocal microscope (Leica SP8). The rate of cell proliferation was calculated by counting the number of EdU-labelled nuclei out of the hundred nuclei observed by DAPI staining.

### Cell cycle inhibitor treatment and expression analysis of KRP3

For cell cycle inhibitor treatment to Tp309 root, 4mg/ml HU, 0.1mg/ml mimosine, 15µg/ml oryzalin and 15µg/ml propyzamide were added into the liquid growth media. After 4h, root samples were harvested for KRP3 expression analysis. Total RNA was isolated using Plant RNA isolation Kit as per the manufacturer’s instruction (Qiagen). cDNA was synthesized using RevertAid H Minus First Strand cDNA Synthesis Kit (Thermo Scientific) using 2µg total RNA. qRT PCR analysis was carried out in Applied Biosystems QuantStudio 3 Real-Time PCR System with three technical replicates and the experiment was repeated with three biological replicates. Two internal control genes Ubq5 and Ef1a were used. Primers used for expression study are mentioned in the supplementary table 1.

### HU sensitivity assay in rice root

For HU sensitivity assay in rice roots, all lines were simultaneously grown in ½ strength MS media containing 0.8% agar and supplemented with or without 2.5mM HU. After seven days, root length was measured for all the lines grown with or without HU. Percent reduction in root length was calculated with the following formula.

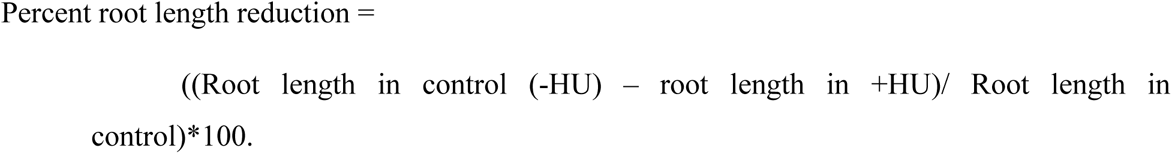

### Yeast Two-Hybrid Assay (Y2H)

Full-length KRP3 coding regions were cloned pGBKT7 vectors. While MAP Kinases (MPK3, MPK4 and MPK6) cloned into pGADT7 have been reported previously by Singh et al., (2024). For yeast transformation, yeast-competent cells of Y2H GOLD strain were prepared according to manufacturer’s instructions (Fast™ Yeast Transformation, G-Biosciences). The two respective constructs were co-transformed in Y2H GOLD competent cells. Co- Co-transformants were initially selected on nutrient medium lacking Leu and Trp (Synthetic Defined (SD)-/Leu/Trp). The resultant co-transformed cells were then streaked on a quadruple drop-out medium deficient in Adenine, Histidine, Leucine and Tryptophan (Synthetic Defined (SD)-/Ade/His/Leu/Trp) followed by incubation at 30°C for 36–48 h. The fully-grown colonies on quadruple drop-out medium were noted as positive interactions.

### In-vitro pull-down assay

For pull-down assay KRP3 was cloned into pGEX-4T-2 vector and transformed into BL21-DE3-Rosetta strain for protein induction. GST-KRP3 protein, induced at 22°C for 12h with 1mM IPTG was immobilized into glutathione Resin (G-Bioscience), as described by Banerjee et al., (2023). Immobilized KRP3 protein was used as prey while bacterially expressed, purified 6xHis-MPK3, 6xHis-MPK4 and 6xHis-MPK6 was used as bait. The binding of KRP3 with MAP Kinases was conducted for 2h at 4°C in binding buffer (50mM HEPES pH 7.5, 150mM NaCl, 2mM EDTA, 5% glycerol, 1% TritonX-100). Post-binding washing was conducted with a binding buffer five times. Finally protein complex was incubated with 5 × Laemmli SDS loading buffer at 98°C for 5 min. For input 20ul of GST-KRP3 bound to glutathione resin and purified MAP kinase (5ug) were used. The presence of prey and bait was detected using western blotting with monoclonal anti-GST and Anti-6xHis antibody (Invitrogen).

### Split Luciferase Complementation (SLC) assay

SLC assay was conducted as described by Chen et al., (2008). In brief KRP3 was cloned into pCAMBIA-CLuc vector while MPK3 was cloned into pCAMBIA-NLuc vector. The clones along with empty vectors were transformed into *A. tumefaciens* strain EHA103. For visualization of interaction, constructs were agro-infiltrated into *N. benthamiana* leaf. Post 48h luciferase activity was detected by CCD camera (ChemiDoc Imaging Systems, Bio-Rad), using a luciferase assay system (Promega).

### Intracellular protein localization

For identification of KRP3 and its phosphor dead and phosphor mimetic forms localization, all three forms of KRP3 were cloned in frame with mCherry at its N-terminal site and 2A-NLS-mGFP .at its C-terminal site. The construct was transiently transformed into *N. benthamiana* leaf. mCherry and GFP fluorescence was visualized 48h post infiltration under a confocal microscope.

### Bimolecular Fluorescence Complementation (BiFC) assay

BiFC assay was carried out as described by Waadt et al., (2008) and Verma et al., (2021). In brif KRP3 was cloned in pSPYNE(R)173 vectors while MPK3 cloned into pSPYCE(M) was used for KRP3 and MPK3 interaction study. While CDKA1 and CycD2 cloned in pSITE-cEYFP-N1 and KRP3 cloned in pSITE-nEYFP-C1 was used for CDKA1/CycD2-KRP3 interaction study. The construct was agro-infiltrated *N. benthamiana* leaf in their respective combinations along with helper construct P19. The leaves were observed 48h post infiltration using a confocal scanning microscope (TCS SP5, Leica) with YFP filters. Excitation and emission wavelengths used were 514 nm and 527 nm respectively.

### *In vitro* kinase assay

KRP3, MPK3 and AtMKK4 were cloned into pGEX-4T-2 vector for GST-tagged protein expression. In-vitro kinase assay was done as described by Jalmi and Sinha, (2016) . Proteins (Kinase and substrate) were incubated with kinase reaction buffer (50mM Tris–HCl, pH 7.5, 1mM DTT, 10mM MgCl_2_, 10mM MnCl_2_, 50mM ATP, and 0.037 MBq of (γ^32^P ATP) [60 Ci/mmol]) at 30°C for 30 min. Kinase reactions were stopped after 30 min by adding 5X SDS loading dye and heated for 5 min at 95°C. Reaction products were run on SDS-PAGE gel and were analyzed by autoradiography.

### Site-directed mutagenesis

The putative phosphorylation target sites (Ser-Pro) were mutated to Ala-Pro or Glu-Pro by base overlap extension PCR. Primers were designed complimentary to each other containing the required mutation. Initially, the target gene was amplified in two parts using gene-specific forward, mutation-specific reverse and mutation-specific forward, gene-specific reverse primer. Both the amplicons were gel purified and used for base overlap extension PCR. The PCR condition for overlap PCR is as follows: 98°C for 30 sec, 55-58°C for 1 min, 72°C for 1 min. After 10 rounds of PCR amplification, gene-specific primers were added and an additional 35 rounds of amplification were carried out. Final PCR products were gel purified and mutations were confirmed using DNA sequencing. After confirmation of mutation amplicons were cloned into desired vector.

### Cell-free degradation assay

KRP3, KRP3^AA^ and KRP3^EE^ with 3’ 6xHis codon were cloned in-frame with MBP tag in pMAL-c2X vector to express and purify MBP-KRP3/KRP3^AA^/KRP3^EE^-6xHis recombinant protein. Total plant protein was isolated by homogenizing Tp309 seedling in protein extraction buffer containing 50 mM HEPES pH 7.5, 150 mM NaCl, 10mM EDTA, 5% glycerol, 5mM DTT, 10mM ATP. 10µg of recombinant KRP3, KRP3^AA^ and KRP3^EE^ protein was incubated with 500µg of plant protein extract for 120 min at 4° C. Reaction was stopped by adding 5xSDS sample loading buffer. KRP3 protein level was detected by western blotting with anti-6xHis antibody (Thermo Fisher).

### Statistical analysis

One-way ANOVA with Tukey’s multiple comparisons test or Dunnett’s multiple comparisons test have been conducted, to determine the significance, in R studio. Significant difference have been represented by Compact Letter Display (CLD) method.

### Accession Numbers

KRP3:-LOC_Os11g40030, MPK3:- LOC_Os03g17700, CDKA1:- LOC_Os03g02680, CycD2:- LOC_Os07g42860

## Author contributions

GB and AKS planned the study and designed the experiments. GB carried out most of the experiments. SJ, UP, BR and DS helped in generating transgenic and recording part of phenotypic data. GB and AKS analyzed the data and wrote the manuscript. AKS approved the final draft.

## Acknowledgements

GB, UP and DS acknowledge the Council of Scientific and Industrial Research, Government of India, and BRIC-NIPGR for fellowship. S.J. acknowledges the Department of Biotechnology, Government of India for fellowship. AKS thanks Sir JC Bose Fellowship from the Science and Engineering Research Board, Department of Science and Technology, Government of India. The authors also acknowledge the Gene Functional Analysis Platform for Crops, Confocal Facility, DNA sequencing facility, Radioisotope facility and the Central Instrumentation Facility of NIPGR, New Delhi, India. The authors declare no conflict of interest.

## Supplemental Data

**Supplementary Figure 1:** Phylogenetic analysis of KRP genes belonging to monocotyledon clade.

**Supplementary Figure 2:** Screening and selection of *krp3* knocked-out plants.

**Supplementary Figure 3:** Screening and selection of KRP3 overexpression transgenic lines and vector control (VC) lines.

**Supplementary Figure 4:** Screening and selection of ProKRP3::GUS and VC-GUS promoter-reporter lines.

**Supplementary Figure 5:** Detection of KRP3 promoter activity using β-glucuronidase (GUS) protein activity assay in elongating internode.

**Supplementary Figure 6:** Activity of KRP3 in developing rice seedlings.

**Supplementary Figure 7:** Detection of KRP3 phosphorylation by MPK3 and identification of specific site of phosphorylation.

**Supplementary Figure 8:** *mpk3* knockout line screening and identification.

**Supplementary Figure 9:** Screening for *krp3mpk3* double mutant lines developed using CRISPR-Cas9 tool.

**Supplementary Figure 10**: Detection of effect for MPK3 mutation in rice vegetative and yield parameters.

**Supplementary Figure 11:** MPK3 mutation moderately affects seedling growth and cell proliferation.

**Supplementary Figure 12:** Selection of KRP3^AA^-OE and KRP3^EE^-OE overexpression line.

**Supplementary Figure 13:** Estimation of vegetative growth parameter of KRP3^AA^-OE and KRP3^EE^-OE lines.

**Supplementary Figure 14:** Comparative estimation of seed number and seed size in KRP3^AA^- OE and KRP3^EE^-OE lines.

**Supplementary Figure 15:** Comparison of shoot and root length of KRP3^AA^-OE and KRP3^EE^- OE with Tp309 and VC seedlings.

**Supplementary Figure 16:** Intracellular localization of KRP3, KRP3^AA^ and KRP3^EE^.

**Supplementary Figure 17:** Phosphorylation does not regulate KRP3’s ability to interact with CDKA1 and CycD2.

**Table S1:** List of primers used in this study

